# Wildfire drives a net decrease in forest live biomass across the Western United States

**DOI:** 10.64898/2026.04.30.720232

**Authors:** Claire M. Zarakas, Grayson Badgley, Michael L. Goulden, James T. Randerson

## Abstract

It remains challenging to quantify recent changes in forest carbon due to lags in forest inventory measurements. The national U.S. forest inventory remeasures plots every five to ten years, so quantifying current carbon stocks using inventory data requires extrapolating from the last time plots were measured. We address this extrapolation challenge by fusing spatially explicit fire disturbance and canopy cover data from Landsat with forest inventory data using a statistical model. We produce annual estimates of live forest carbon across the Western U.S. from 2005 to 2022, and find that live forest biomass increased from 2005 to 2015, and then declined by 5% from 2015 to 2022 — a signal missed by both official U.S. reporting and Earth system models. The trend reversal was driven primarily by increasing tree mortality from wildfire, and secondarily by slowing rates of carbon accumulation in undisturbed areas. Our results highlight the importance of accounting for rapidly changing disturbance regimes, and can help to improve jurisdictional carbon accounting and inform the extent to which federal and state climate mitigation strategies can rely on land to achieve net-zero emissions targets.

**Significance statement:** Policy makers need to accurately and rapidly assess the status of the land carbon sink in order to make land management decisions and to assess progress towards climate commitments. However, lags in on-the-ground measurements make it challenging to do so, and it remains an open question whether Western U.S. forests are a net sink or a source of carbon. We fuse on-the-ground forest measurements with remote sensing data to show that live biomass is net declining in Western U.S. forests, and that this trend is driven primarily by increasing wildfire activity. This result challenges the idea that jurisdictions can rely on the land to offset fossil emissions, and supports tracking land carbon trends separately from fossil emissions inventories.

## 1 Introduction

Wildfire exerts a fundamental control on the amount of carbon forests can store, and it is rapidly changing across the Western United States. Burned area has increased approximately tenfold since 1985 (1), individual wildfires have become larger and more severe (1–3), and the 2020 and 2021 wildfire seasons in the West shattered previous records (4). This intensifying wildfire regime has unfolded amid a suite of additional forest stressors, including unprecedented drought (5–8), insect disturbance (9–11), and higher regional temperatures (12, 13). While Western U.S. forests have long been considered a stable carbon sink, the compounding effects of these stressors raises the question: is live biomass still accumulating, or has the intensification of disturbance pushed Western forests into biomass decline?

In principle, it should be straightforward to answer this question by directly measuring trees across the Western U.S. However, the combination of lags in field observations and rapid ecosystem change makes this challenging in practice. The U.S. Forest Inventory and Analysis (FIA) program maintains a national network of forest inventory plots that are measured on a 5- to 10-year cycle. This authoritative dataset underpins official U.S. forest carbon accounting, which informs parts of the U.S. greenhouse gas inventory reported to the United Nations (14). The FIA remeasurement cadence works well for detecting subtle changes in an otherwise slowly changing system, but introduces lags when forest ecosystems are changing faster than plots are remeasured. In fact, only about 15% of FIA plots in the Western United States have been measured since 2020, meaning that the effects of particularly extreme wildfire seasons in 2020 and 2021 (Figure 1) have not yet been fully captured in on-the-ground measurements.

**Figure 1:**
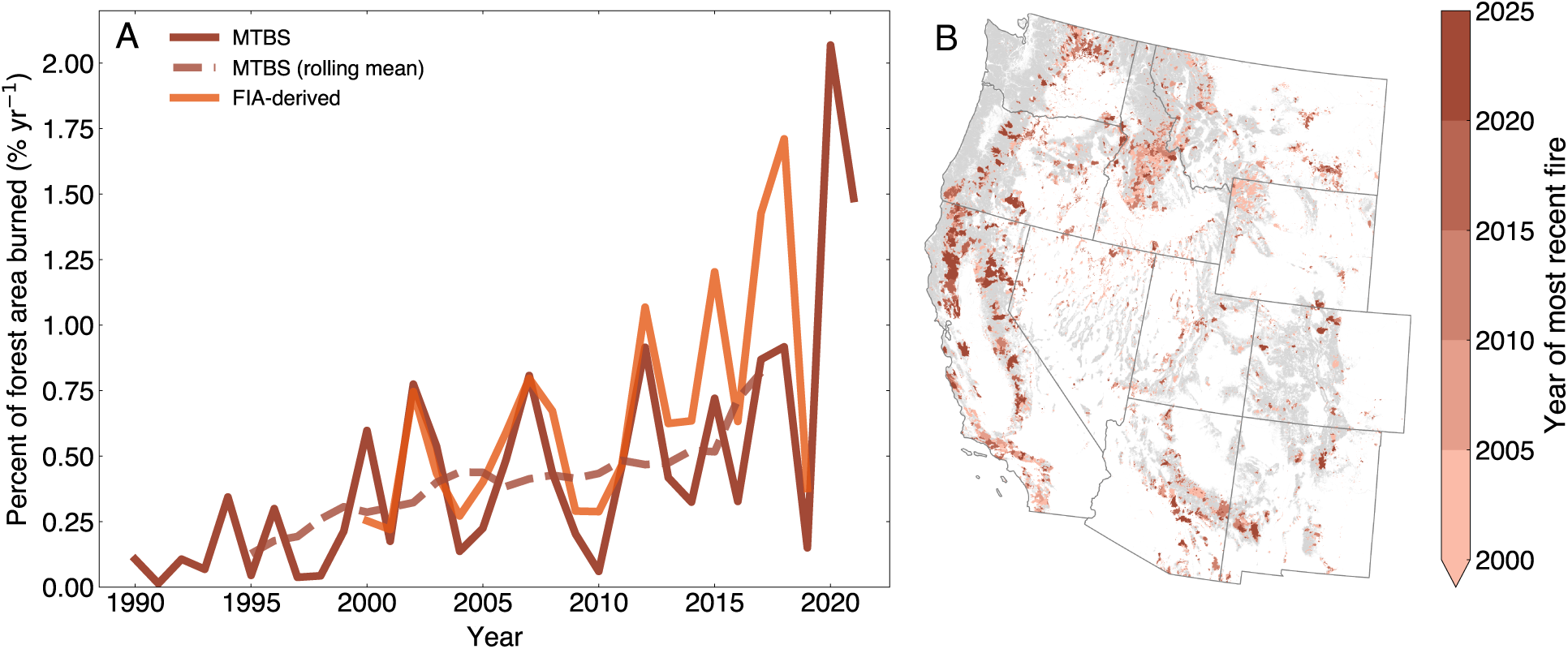
Forest fire trends across the Western United States. (A) Percent of forest area burned annually, calculated from Monitoring Trends in Burn Severity product (21) fire perimeters, and fire disturbance codes recorded in the Forest Inventory and Analysis (FIA) National Forest Inventory (23). We calculate FIA percent burn by dividing the number of plots with a fire disturbance code in each year by the number of all plots measured in or after that fire year. Fires in (A) and (B) are filtered to grid cells that are at least 50% forests according to the Landscape Change Monitoring System land use product (24), which is shown in gray.

These lags limit both the scientific community’s understanding of the carbon cycle and jurisdictions’ ability to track progress towards climate commitments and ecosystem management goals. Many states have incorporated forest carbon sinks into their climate targets, so forest carbon trends directly inform economy-wide climate mitigation strategies. Similarly, land managers require up-to-date information about the state and trajectory of the forests they oversee to guide management decisions, like when to suppress wildfires and where to prioritize restoration efforts.

Several studies have estimated recent live biomass trends across the Western United States and broader domains using inventory data (14–18), remote sensing (19), and process-based models (20). However, these efforts have not converged on a consistent answer to how regional biomass stocks might be changing. Part of this disagreement relates to how each approach navigates trade-offs between incorporating plot-level field measurements and the need to better capture the effects of changing disturbance rates. Despite rapid intensification of disturbance across the Western United States, the fundamental question of whether forests are net accumulating or losing biomass has not been directly answered.

Here, we attempt to answer this question by fusing forest inventory data with satellite-derived disturbance data in a statistical approach that is both grounded in plot-level field measurements and capable of capturing recent disturbances (see Methods). We produce annual live biomass estimates from 2005 to 2022 by combining three random forest models that are trained on FIA plot-level data across the Western United States (see Figure S1). These models capture observed relationships between biomass and fire occurrence, meteorology, canopy cover, and other predictors. The first model estimates initial live aboveground biomass stocks in 2005 (Table S1). The second model estimates annual changes in aboveground biomass in burned areas. The third model estimates annual changes in aboveground biomass in unburned areas, capturing both accumulation in undisturbed forests and the effects of non-fire disturbance, such as harvest and drought. We apply these models forward in time using spatially continuous inputs, including annual wildfire perimeters from Monitoring Trends in Burn Severity (MTBS, ref. 21) to capture year-to-year changes in fire disturbance (Figure 1). Training each model on FIA plot measurements, while forcing them with annually updated and spatially continuous data derived from remote sensing, grounds our approach in field observations while also incorporating time-varying disturbance rates.

We find that live biomass in Western U.S. forests increased through 2015 and then declined by 5.1% through 2022, driven primarily by wildfire-induced tree mortality. We compare our estimates against a range of live biomass estimates spanning national to global scales, including official U.S. carbon accounting (14, 22), FIA-based academic studies (15–18), satellite-derived estimates (19), a process-based land model (20), and CMIP6 Earth system models. The declines we document are broadly consistent across several independent lines of evidence. However, the official U.S. carbon stock estimates and CMIP6 Earth system models show the opposite trend, indicating live biomass increases. This divergence likely stems from a common cause: both approaches underestimate the rate at which disturbance has intensified in recent years.

## 2 Results

### 2.1 Model evaluation

We evaluate our model by comparing it to FIA data during the periods when plots have been measured at least twice. In the Western United States, this roughly corresponds to 2005 to 2016, because on average, plots were first measured in 2005 and then re-measured in 2016. Over this period, our model captures broad spatial patterns of changes in live biomass stocks at the 0.25° spatial scale (R² = 0.55, see Figure S2 and SI Section S1.3). Coastal forests showed net carbon accumulation, indicating that background productivity rates have outpaced losses from disturbance, even though there are high rates of fire and harvest in these high biomass regions (25). Biomass declined in parts of the Mountain West, particularly Colorado, central Idaho, and western Montana. The spatial distribution of these losses is consistent with previously documented patterns of bark beetle outbreaks that occurred from approximately 2005 to 2012 (9–11). However, our model underestimates live biomass losses in these mountain regions prior to 2015, likely due to our model lumping all unburned plots together and not accounting for differences between different non-fire disturbance types (SI Section S1).

### 2.2 Trends in forest live aboveground biomass

While we can only directly compare our estimates to FIA data through 2016 on average, we estimate regional live aboveground biomass for the period from 2005 through 2022 by assimilating burned area and canopy cover data derived from Landsat and climate predictors from the Parameter-elevation Regressions on Independent Slopes Model (PRISM, 26). We find that live aboveground biomass in forests across the Western United States decreased from 2005 to 2022 (Figure 2A).

**Figure 2:**
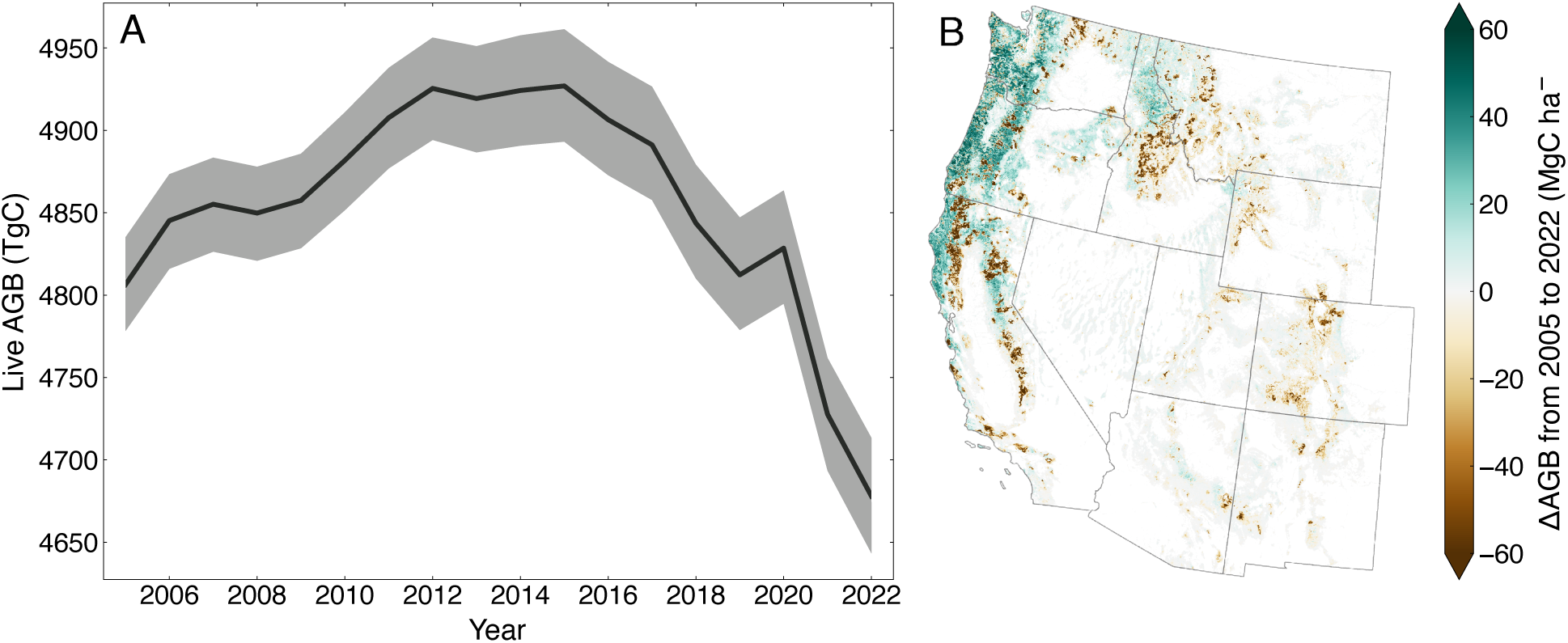
Change in live forest aboveground biomass carbon from 2005 to 2022 across the Western United States, shown as (A) annual estimates of Western U.S. forest aboveground biomass, in Teragrams of carbon (TgC) and (B) the cumulative change in forest live aboveground biomass density (MgC ha^−^¹) from 2005 to 2022. Uncertainty bounds in (A) reflect the 5th to 95th percentile range from a 500-member ensemble.

Using a piecewise linear regression, we identify the turning point between net carbon accumulation and net carbon loss in 2015 (p<0.05). The rate of post-2015 decline is especially notable when contrasted against the relatively slow rates of net carbon gain from 2005 to 2015. Live biomass stocks increased by 14 TgC yr^−^¹ from 2005 to 2015 (a 2.9% increase overall), and then reversed, declining by 37 TgC yr^−^¹ from 2015 onward (Figure 2A). By 2022, the Western U.S. carbon stocks fell below 2005 levels, a 2.7% (5th–95th percentile range 2.0 to 3.3%) decline from 2005 and a 5.1% (4.8 to 5.3%) decline from their 2015 peak. Apart from a single year (2019), the decline has been relatively steady and unidirectional. These trends are robust when we look over an ensemble of 500 simulations, each of which samples a subset of the data (light grey in Figure 2A, see Methods). Cumulative regional biomass losses from 2015 to 2022 are equivalent to approximately 15% of fossil CO₂ emissions from Western U.S. states over the same period (Table S2).

Live biomass losses were not evenly distributed across space (Figure 2B). The largest declines occurred in the northern Sierra Nevada, and the largest losses occurred within the perimeters of recent fires (Figure 1B). Meanwhile, coastal California and wet forests west of the Cascades in Oregon and Washington saw almost uniform gains. For drier, interior forests, losses were concentrated in central Idaho and across most of Colorado and New Mexico. The regional decline in live biomass stemmed from substantial losses in just 22% of the Western U.S. forests, which were large enough to counteract the live biomass gains that occurred over 32% of the West, and minimal changes over the rest of the domain (Figure 2B, Table S3). The spatial pattern of biomass losses (Figure 2B) mirrored the mosaic of fire perimeters in Figure 1B. 2020 and 2021 account for the largest share of post-2015 biomass loss (Figure 2A), reflecting especially active fire years (Figure 1A).

### 2.3 Disentangling drivers

Because our model tracks wildfire-driven carbon stock changes separately from net growth in unburned areas, we can quantify how much wildfire contributed to the shift in trends in biomass between 2005 to 2015 and 2015 to 2022 (Figure 3). Wildfire caused 77% of the change, while decreasing accumulation rates in unburned areas explain the remaining 23% (Table S4). Across the Western United States, losses from wildfire were primarily driven by increases in burned area (Figure 1A-B), as opposed to increases in rates of tree mortality. On average, our model estimated that wildfires killed 43% of live aboveground biomass, which remains approximately constant over time (Figure S3), and is consistent with the mean 40% loss observed in forest inventory plots (Figure S3, see also ref. 27). Most forests with large fractional carbon losses from 2005 to 2022 were areas that burned at some point during that period (Figure 3C).

**Figure 3:**
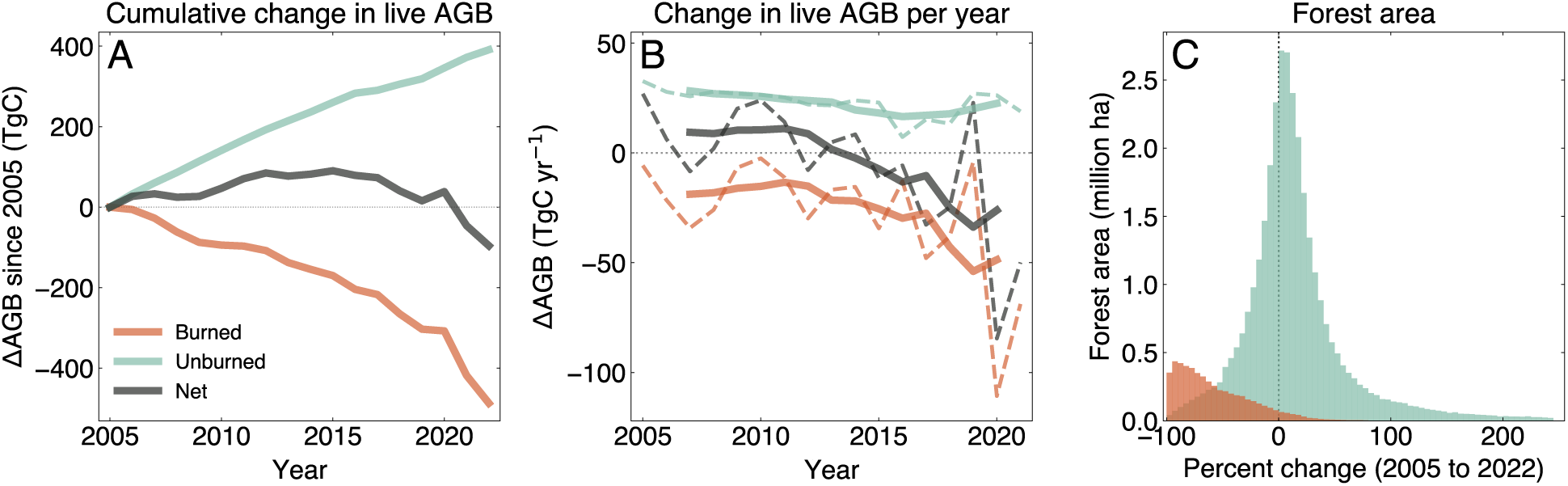
Drivers of changes in live forest biomass. (A) Annual changes in live aboveground biomass (AGB) across the Western United States due to fire (red) and carbon changes in unburned areas (teal), shown both for each year (dashed) and as a centered 5-year rolling average (solid). (B) Cumulative changes in live AGB, partitioned into the effects of fire and carbon changes in unburned areas. In (A) and (B), we only count forests as “burned” during the year that a fire occurred — forests that burned in prior years are included in the “unburned” signal, so post-fire regrowth contributes to the unburned. (C) Distribution of percent cumulative changes in live AGB carbon from 2005 to 2022, in any grid cell that burned over that time period (red) compared to grid cells that experienced no fires during the study period (teal)

In unburned forests across the Western United States, net biomass accumulation rates declined from 2015 to 2018, before recovering after 2019 (Figure 3A, teal line). These net biomass accumulation rates reflect the effects of both tree growth and biomass losses from non-fire disturbances. Our approach does not quantitatively disentangle the contribution of these two effects in undisturbed areas, but the spatial pattern (Figure S4) aligns with patterns of drought-driven forest mortality in California (28), which were concentrated in the southern Sierra Nevada. The timing is also consistent (Figure S4E), because while the drought occurred from 2012 to 2015, most mortality occurred in 2015 and 2016 (28, 29). The post-2019 recovery in unburned forest accumulation rates likely reflects both a return to lower drought-driven mortality rates and contributions of post-fire regrowth in previously burned forests.

### 2.4 Comparison with other Western U.S. estimates

Several recent studies support our results about the general pattern of live biomass decline across Western U.S. forests (Figure 4). Li et al. (19) examined Northern Hemisphere biomass trends using radar-derived estimates of vegetation optical depth and found modest gains in live Western U.S. forest biomass prior to 2016, followed by sustained losses through 2022. The timing and sign of that decline closely mirrors the live biomass reversal we detected using an independent line of evidence. Similarly, Liu et al. (20) used a process-based vegetation model and found a net decline in Western U.S. live forest biomass from 2005 to 2020. While their model exhibits larger interannual variability and does not identify a clear sink-to-source transition year, their overall declining trend is consistent with our results.

**Figure 4:**
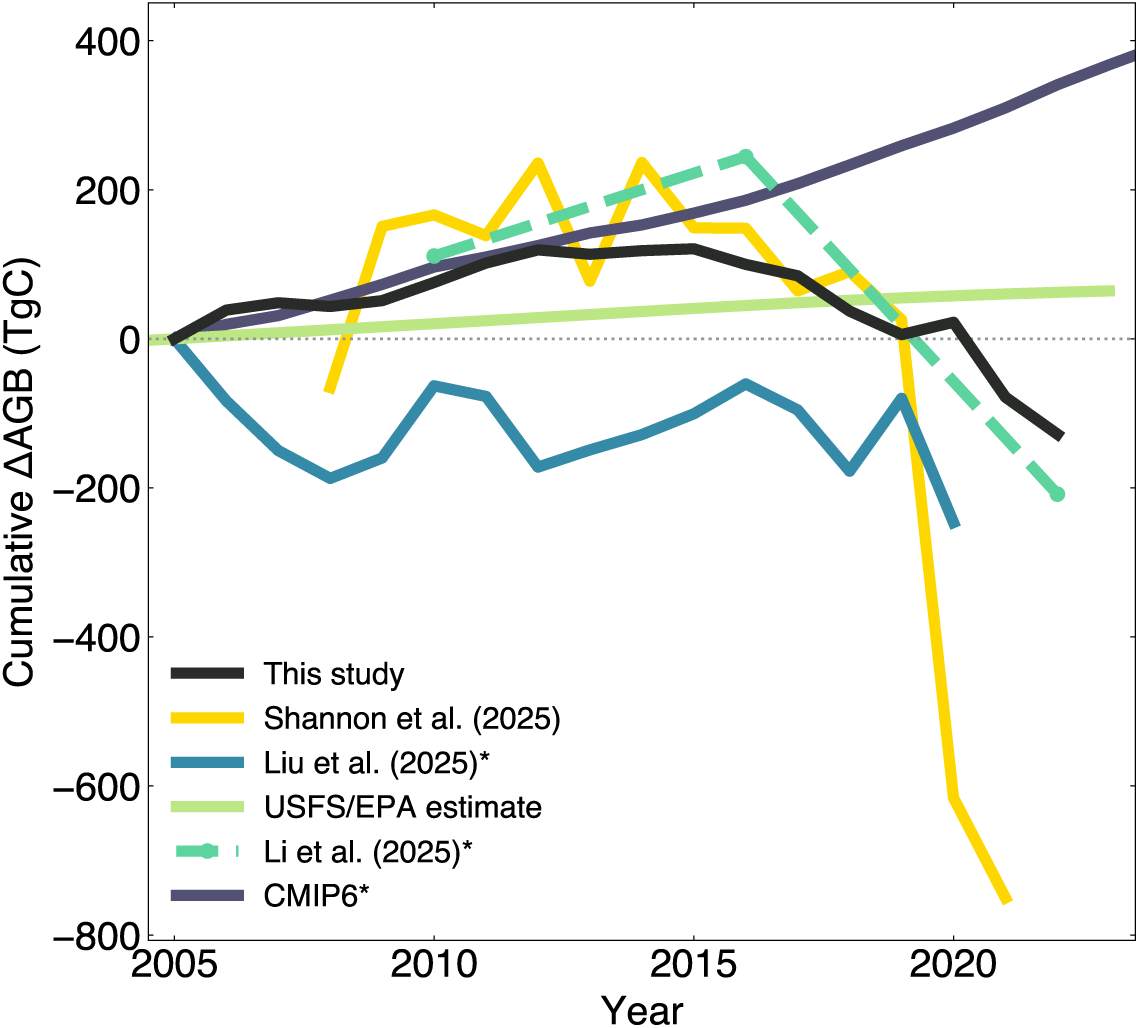
Comparison to trends in other biomass products. Starred datasets (Li et al. (19), Liu et al. (20), and CMIP6) quantify total live biomass, including both forest and nonforest carbon. We plot total live biomass carbon changes from this dataset, because the majority of carbon changes in the Western United States are driven by forest carbon changes. Starred datasets are normalized by the percent of forest remaining forest in Figure S5. Because the Li et al. (19) study spanned 2010 to 2022, we plotted 2010 based on a linear extrapolation from 2005, in order to facilitate comparison of cumulative change in live biomass since 2005. Li et al. (19) is shown as a dashed line because they report trends during two time periods rather than annual biomass estimates. CMIP6 biomass estimates were calculated as total vegetation carbon (‘cVeg’) multiplied by a factor of 0.8 to convert total live biomass to aboveground live biomass (following ref. 30), and the multi-model mean is shown here.

Our results are also broadly consistent with studies based on FIA data (15–18). Hall et al. (16) found live biomass declines in much of the Western United States from 2005 through 2019, largely located in Intermountain West. Shannon et al. (17, 18) studied a longer period and found statistically significant declines primarily in the Sierra Nevada, with large declines in 2020 and 2021 (Figure 4). Meanwhile, Hogan et al. (15) found neutral or declining forest productivity across all Western U.S. ecoprovinces, even after accounting for disturbance-driven mortality. While these studies generally suggest a decline in Western U.S. forest biomass, they disagree as to the magnitude and location of those declines (15–18). Despite all relying on the same core dataset, it is still challenging to compare FIA-based studies to each other — or to other methods, like process-based models — because these studies do not produce gridded annual estimates of biomass stocks, and each focuses on different time intervals.

Notably, both the official U.S. greenhouse gas inventory and estimates from Earth system models (ESMs) participating in the Coupled Model Intercomparison Project version 6 (CMIP6) show trends in live biomass that are opposite in sign to our results. These approaches estimate that live biomass has increased by 1 and 4%, respectively, from 2005 to 2022, compared to our study’s 3% decline. These trends likely both diverge from our study for the same reason: underestimation of disturbance rates.

The United States Forest Service (USFS) and United States Environmental Protection Agency (EPA) estimate land carbon stocks for use in the U.S. national greenhouse gas inventory program (hereinafter referred to as the official U.S. approach, Figure 4, lime green line) using FIA-derived transition rates between different stand age classes and relationships between stand age and biomass density. This approach results in a near-linear increasing trend in live biomass across the Western United States (Figure 4), driven by gains in California, Oregon, and Washington that offset declines in inland states. Both the official U.S. approach and our approach rely on the same underlying national forest inventory dataset. During the period where no extrapolation takes place, these approaches largely agree, both in terms of absolute carbon stocks (Figure S6) and the sign of trends in carbon stocks (Figure S7). However, the trends diverge in sign once we move beyond the well-observed period, which traces back to how the two approaches account for changing disturbance rates.

The official U.S. approach implicitly assumes that disturbance rates observed over the historical record remain approximately constant when extrapolating forward in time (14, Annex 3 Part B). Our approach, however, accounts for changing disturbance rates by incorporating satellite-derived estimates of both burned area and changes in canopy cover. These small changes — allowing fire disturbance rates and canopy cover to update through time based on observations — results in a significant divergence between the two approaches. From 2015 to 2022, the official U.S. approach reports live biomass stocks increasing by 1%, while our approach finds they decline by 5%. Eventually, burned plots will be remeasured, and those losses will be incorporated into estimates from the official U.S. approach. But until then, and as fire activity further intensifies, long intervals between plot remeasurement will likely result in the systematic overestimation of live biomass stocks under the official U.S. approach.

Our results likely diverge from the CMIP6 ensemble for similar reasons. It is widely acknowledged that the land component of ESMs have limited (if any) representation of tree mortality processes (31), including both fire- and drought-driven mortality (32, 33). For example, both ESMs and models used in the “Trends and drivers of the regional scale terrestrial sources and sinks of carbon dioxide” (TRENDY) project struggle to accurately simulate historical burned area (34), which matters for historical simulations because fire occurrence is dynamically calculated rather than prescribed by observations. Further, ESMs have simplistic representations of how drought affects the terrestrial carbon cycle (35). Underestimating disturbance rates, and how those disturbance rates change over time, can result in the overestimation of live biomass stocks by ESMs, which closely mirrors the dynamic we identified in the official U.S. approach.

## 3 Discussion

### 3.1 Opportunities to nowcast terrestrial carbon

Adaptive management of terrestrial carbon stocks requires that carbon inventories be both accurate and up to date. Land managers rely on rapidly updating information in order to decide how to resist, accept, or direct forests in transition (36), and these decisions should be frequently revisited to assess whether they are having the desired impact on ecosystems (37, 38). This feedback cycle requires carbon stock estimates that update as fast as ecosystems are changing.

Carbon tracking systems based on on-the-ground forest inventory observations alone are unable to meet this need. In the Western United States, the current rate of forest inventory plot remeasurement is not keeping pace with rates of change on the ground. The limitations of inventory-only approaches are evidenced by the divergence between our approach and the official U.S. approach for estimating carbon stocks (Figure 2B). Remote sensing data can bridge these temporal gaps. By integrating remote sensing and inventory data, we demonstrate the beginnings of an approach that captures changes that have already happened but are not yet reflected in field measurements, enabling carbon stocks to be more accurately extrapolated forward to estimate current forest conditions.

Fully implementing a carbon tracking system that fuses forest inventory and remote sensing data operationally and at scale would require further development, and has been a longstanding goal of terrestrial carbon cycle science (17, 39–41). Several extensions to our approach would be particularly critical for achieving this goal. First, the approach would need to encompass all ecosystem carbon pools — not just live biomass — in a way that ensures physically realistic, mass-conserving fluxes between pools. Second, explicitly modeling non-fire disturbances, particularly harvest and insect disturbances, would improve estimation of biomass losses. This extension would require continuously updated, accurate wall-to-wall annual disturbance maps, and could potentially build on recently developed gridded disturbance products (25). Finally, operationalization requires sustained investment in both the annual gridded products the approach would depend on, and in the FIA national forest inventory program itself. Approaches that incorporate remote sensing data do not replace the need for rigorous, on-the-ground forest inventories: inventory data remain the gold standard for detecting long-term changes, and for training statistical or mechanistic models that assimilate remote sensing data.

### 3.2 Fate of the Western U.S. carbon balance

We show that Western U.S. live biomass stocks, long considered a carbon sink, have become a carbon source. Because we only model live biomass, however, we cannot assess whether Western U.S. forests have fully transitioned from being a carbon sink to a source. That determination requires quantifying total ecosystem carbon storage, including dead biomass, soil, and litter carbon pools. We note that our main result is not a projection of future risk — it represents our estimate of biomass that has already died but has not yet “hit the books” due to lags in the remeasurement of FIA plots.

While this transition has likely not yet fully propagated to the forests’ total ecosystem storage, the ecosystem-level shift from a sink to a source is likely locked in. Many fire-killed trees remain standing, and will fall during the next 10 years and decay in the following decades (42), releasing carbon that is effectively committed to the atmosphere even if the actual emissions have not yet occurred. This live biomass sink-to-source transition is also consistent with forests adjusting to a new, lower equilibrium carbon density under a warmer climate (13, 43, 44).

Looking forward, there is little sign that this trajectory will reverse in the near term, though it is possible fire-fuel feedbacks could eventually constrain burned area over the longer term (45). Roughly 23% of California’s arid forests have burned in the past ten years, compared to just 7% of arid forests in Washington and Oregon (Figure S8). In other words, there are still plenty of dry forests in the Western U.S. that have yet to burn. And with fire activity projected to continue to intensify across the Western U.S. (45, 46), carbon losses from future fires will likely outpace regrowth in previously burned forests. While trees will regrow in post-fire landscapes, recovery can take decades (47, 48), and prior work suggests forests will recover to a lower carbon density than before due to anticipated shifts in ecosystem composition (13, 43).

We focused on the Western U.S. because it has already experienced well-documented and rapid changes in disturbance regimes. However, the intensification of forest disturbances is a global phenomenon (49–51). Our study adds to a growing body of work documenting a weakening global land carbon sink (52) and even sink-to-source transitions in some regions (53–56). Our results are also consistent with recent work identifying a discrepancy between remote-sensing based live biomass estimates and atmospheric inversions (30, 57) that suggest a weaker global live terrestrial biomass carbon sink than official inventories currently reflect.

### 3.3 Implications for climate commitments

Finally, our study demonstrates the risks of overrelying on terrestrial carbon storage for addressing climate change. Most climate commitments target net-zero emissions or a specified percent reduction in net emissions, meaning that land carbon estimates directly shape expectations for ambition around other mitigation tools, such as direct fossil fuel emissions reductions. Many federal and state plans to achieve net-zero emissions explicitly assume a positive land sink that offsets a portion of fossil fuel emissions (58, 59). The U.S. long-term climate mitigation strategy assumes the land sink will remain roughly constant through 2050 (60). Globally, an analysis of 2030 National Determined Contributions under the Paris Agreement found that land-based carbon removals accounted for nearly 25% of all pledged mitigation activity (1.5 ± 1.1 GtC yr^−^¹, ref. 61). Accurate, up-to-date estimates of the current status and recent changes in terrestrial carbon stocks can help align these political goals with the realities of the global carbon cycle (62–64). A weaker land sink (or even a source, as we show in this study) would imply the need for more ambitious climate action in other sectors to achieve the same climate targets.

Our findings also reinforce growing calls to track fossil fuel emissions and land carbon fluxes separately — on a like-for-like basis — in both accounting frameworks and climate targets (65–67). Counting all of the land sink toward net-zero targets is physically inconsistent with the goal of stopping anthropogenic warming, because allowing the passive component of the land sink to count toward net-zero goals would result in global temperatures continuing to rise even under nominal net-zero emissions (67). Tracking fossil fuel emissions progress separately can also guard against climate mitigation plans that rely on sustained land uptake, only to have the land sink weaken under climate change. Further, separating fossil fuels and land carbon enables more effective tracking of the impact of climate policies. Fossil fuel emissions are under direct human control and can be reduced by climate mitigation policy.

Land carbon fluxes, on the other hand, reflect a combination of ecosystem demographic changes, land management decisions, and the effects of climate change. Tracking land and fossil emissions separately both better isolates the impact of climate policy, and can help guard against overreliance on land in mitigation strategies.

## 4 Methods

We calculate annual live forest biomass carbon across the Western United States by combining plot-level biomass data with spatially explicit meteorological and disturbance data. First, we use plot-level FIA biomass data to train a statistical model of how disturbance and climate affect carbon accumulation rates over time. Then we run our model forward and backward from 2005 using spatially explicit fire disturbance and canopy cover data. We limit our analysis to “forest remaining forest” in order to isolate forest demographic changes, as opposed to land use transitions. We adopt the U.S. Forest Service definition of forestland, which is defined as an area that has at least 10% canopy cover of trees, or has had at least 10% canopy cover that can be naturally or artificially regenerated. As such, our analysis includes the effects of human and natural forest disturbance (e.g., fire, harvest, etc.), but does not include the effects of permanent deforestation (e.g., forest conversion to cropland) or afforestation.

### 4.1 Model structure

We estimated biomass with a statistical model driven by plot-level FIA data. Our model consisted of a combination of three random forests: one initial biomass estimator (Section 4.1.1), and two biomass change estimators, conditioned on fire occurrence (Section 4.1.2). Each random forest contained 300 trees with a maximum tree depth of 30. We estimated uncertainty by training 500 different sets of models using 500 random subsets of 80% of the FIA training plots. We used SHapley Additive exPlanations (SHAP) values (68) to quantify the importance of each predictor in each model component.

We developed separate estimators for absolute biomass and for changes in biomass, to disentangle the drivers of spatial and temporal variability in biomass stocks. For example, we expected mean climate to be the primary driver of broad spatial patterns of biomass stocks, but expected for climate variability to exert a smaller control on changes in biomass at any given location over time. Constructing separate estimators for spatial and temporal variability allowed us to better represent how the same variables can exert different mechanistic controls on spatial versus temporal biomass patterns. We use two biomass delta estimators: one for carbon stock changes in plots that burned (Section 4.1.2.1), and one for carbon stock changes in plots that were not burned (Section 4.1.2.2). Each of these biomass delta estimators use the biomass from the previous timestep as a predictor, so the full model runs recursively forward over time using spatially continuous gridded input data (see Section S1.1 and Section S1.2.2).

#### 4.1.1 Carbon stock initialization

We estimate regional baseline carbon stocks in 2005 by training a random forest model that predicts live aboveground biomass using FIA data from the 2000 to 2009 measurement interval combined with spatially varying meteorological, topographic, disturbance, canopy cover, and geographical predictors. Meteorological predictors included the 10-year mean temperature, maximum warm season temperature, minimum cold season temperature, and mean precipitation. Meteorological variables were calculated from the 10 years prior to the FIA aboveground biomass measurement, based on gridded PRISM data extracted at the fuzzed plot coordinates. Topographic predictors were slope, elevation, and aspect, as recorded in FIA plot-level data. For disturbance predictors, we included years since fire, stand age, and percent public ownership from the plot-level FIA data in the year that the biomass stock was recorded. Finally, we used latitude, longitude, ecoregion and ecoprovince as recorded in FIA data. Stand age and canopy cover were the strongest predictors of initial biomass (Figure S9A), with higher biomass associated with older forests with higher canopy cover (Figure S10A).

#### 4.1.2 Carbon delta models

We model how different drivers influence carbon stock changes over time from forested FIA plots with at least two biomass measurements. We model carbon stock changes as a function of (1) static site characteristics, (2) carbon stocks in the prior measurement, (3) and temporally varying disturbance, meteorology, and canopy cover data.

##### 4.1.2.1 Fire carbon change estimator

We trained a random forest model to predict changes in plot carbon stocks following a fire. To train the model, we used 2,912 FIA plots across Western U.S. forests that had been both i) measured prior to a fire and ii) remeasured following that fire. These paired measurements allowed us to calculate changes in plot biomass from fire, and for our model to infer fire-induced mortality rates. In addition to FIA plot data, the model predicts change in live aboveground biomass carbon based on pre-fire carbon stocks, changes in the National Land Cover Database (NLCD) canopy cover between measurement years, static topographic and ownership predictors, and the mean meteorology over the 10 previous years. Pre-fire carbon stocks and changes in NLCD canopy cover between measurement years provided the most predictive power (Figure S9B), followed by mean meteorology. We assumed that when live biomass losses occured between measurements, all biomass losses occurred in the year that the fire was recorded. When live biomass gains occurred between measurements, we linearly interpolated between measurements to estimate the carbon gain in the fire year (i.e., if a plot gains 10 MgC ha^−^¹ from 2000 and 2010 and a fire occurred in 2005, we assume that the plot gained 1 MgC ha^−^¹ yr^−^¹ every year including in 2005).

##### 4.1.2.2 Unburned carbon change estimator

The unburned carbon change random forest model captures all biomass changes in unburned plots, including the effects of non-fire disturbances like harvest and insect herbivory. Ideally, we would represent each of these disturbance agents with its own separate model, which would allow the relationship between canopy cover change and biomass change to vary by disturbance type rather than assuming a single universal relationship. However, we found that fire was the only disturbance agent that has been reliably mapped at the regional scale, so we combined all non-fire disturbance types into this single model. Our model relied on changes in annual canopy cover to capture these non-fire forest disturbances.

We trained the unburned carbon change estimator on all 36,603 plots that had at least two biomass measurements and that did not experience a fire between measurements. We linearly interpolated between measurements to estimate the unburned carbon change per year. The model also included the following inputs: initial carbon stocks, changes in NLCD canopy cover per year between measurement years, static topographic and ownership predictors, years since fire, and the mean meteorology over the 10 years prior to the second biomass measurement. Similar to the fire carbon change estimator, initial carbon stocks and changes in NLCD canopy cover per year contributed considerable predictive power to the model (Figure S9C). Critically, including initial biomass as a predictor in undisturbed forests allows our model to capture how forest structure influences forest productivity. For example, in a given climate regime, forests with lower initial biomass (i.e., younger forests) grow more each year than forests with higher initial biomass (Figure S10M). Our estimator also implicitly captures how harvest rates vary with forest structure and ownership type — privately owned, high biomass forests are more likely to have biomass declines in a given year than publicly owned high biomass forests.

#### 4.1.3 Comparison to other biomass estimates

We compared our results to several other gridded biomass datasets, including one dataset derived from remote sensing (Li et al, ref 19) and one derived from a process-based model that assimilates burned area (Liu et al; ref 20). These datasets report total live aboveground biomass, including both forest and nonforest carbon. This causes a slight discrepancy in our comparison, because our study only estimates total live aboveground biomass for forests. We compare our results to these studies’ total biomass estimates (main text) because the majority of carbon changes in the American West are driven by forest carbon changes. We also normalize these studies by percent forest cover in the SI. Because the Li et al. study spanned 2010-2022, we plotted 2010 based on a linear extrapolation from 2005, in order to facilitate comparison of cumulative change in live biomass since 2005.

We also compared our results to total vegetation carbon (‘cVeg’) as modeled by CMIP6 models. We converted total vegetation carbon to aboveground vegetation carbon using a factor of 0.8 to convert total live biomass to aboveground live biomass (following ref. 30). We used a combination of the historical scenario and shared socioeconomic pathway 2 – 4.5 (SSP245), which was necessary because the CMIP6 historical scenario ends in 2015. We included all models and ensemble members which had cVeg data available for both the historical and SSP245 scenarios (Table S5).

### 4.2 Model evaluation

We evaluated each random forest separately using 20% of plots withheld from the training dataset for each ensemble member. We evaluated our model at the regional scale by coarsening our dataset to 0.25 degree resolution and averaging biomass estimates across the testing plots in those grid cells. We also report model evaluation at the plot-level (Section S1.3), where our model explains a smaller fraction of observed variance. The poorer performance at the plot level is expected because the fuzzed FIA plot locations prevent us from comparing our model estimates at the exact same location where plots were measured, and there is a high degree of small-scale spatial variability in biomass stock changes.

## 5 Open Research

The input data for our analysis is all publicly available following the references cited in the main text. Processed inputs on a consistent 1,000 meter grid, the processed FIA plot-level dataset, and the outputs of our analysis are available on Zenodo (69).

The code used to produce this manuscript is available on GitHub. All analyses were performed in Python using open source packages (Section S1.4).

## Conflict of Interest

JTR serves on the technical advisory board for the Symbiosis Coalition. The authors declare no other conflicts, financial or otherwise, that influenced, or could be perceived as influencing, the research described here.

## Acknowledgements

CMZ was supported by Schmidt Science Fellows, in partnership with the Rhodes Trust. JTR acknowledges funding support from the U.S. Dept of Energy Office of Science RUBISCO Science Focus Area, the U.S. National Science Foundation Collaborations in Artificial Intelligence and Geosciences program (RISE-2425932), and NASA’s Earth Information System Fire Project, Fire Sense Technology Program (80NSSC24K1823), Fire Sense Implementation Team (80NSSC24K1317), and Carbon Monitoring System Program (80NSSC25K7211). We thank Olaf Kuegler for providing information about the structure of the FIA database. We thank all of the USFS personnel who contributed to the development of the FIA database, including scientists, field crews, and administrators.

## Supporting Information

### S1 Extended methods

#### S1.1 Model structure

Our model estimates changes in live aboveground biomass (AGB) over time by combining three random forest models. We developed separate estimators for absolute biomass and for changes in biomass in order to disentangle different drivers of spatial versus temporal variability in biomass stocks. For example, we expected mean climate to be the primary driver of broad spatial patterns of biomass stocks, but expected for climate variability to exert a smaller control on biomass changes over time at any given location. We use two biomass delta estimators: one for carbon stock changes in plots that burned (Section 4.1.2.1), and one for carbon stock changes in plots that did not burn (Section 4.1.2.2). Each of these biomass change estimators use the biomass from the previous timestep as a predictor, so the full model can be recursively run forward over time (see Section S1.1).

At the initial time step *t* = 2005, biomass is estimated as:

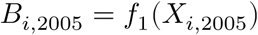

At each time step *t*, biomass is updated based on whether the cell burned:

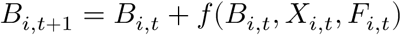

Where

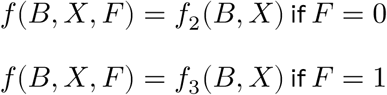

*B*_*i*,*t*_: Biomass in cell

*E*_*i*,*t*_: Exogenous variables (climate, topography, etc.)

*F*_*i*,*t*_: Fire indicator (1 if burned, 0 if unburned)

*f*_1_: Initial biomass estimator

*f*_2_: Biomass change estimator for unburned areas

*f*_3_: Biomass change estimator for burned areas

By assimilating changes in live canopy cover, our model attempts to capture the effects of tree mortality from non-fire disturbance. However, by lumping all non-fire disturbances together, it essentially fits a single relationship between live canopy cover change and biomass change, without distinguishing whether the canopy cover change was caused by insect disturbance, drought, harvest, windthrow, or other disturbance agents. Ideally, we would represent each of these disturbance agents with its own separate model, which would allow the relationship between canopy cover change and biomass change to vary by disturbance type rather than assuming a single universal relationship. However, we found that fire was the only disturbance agent that has been reliably mapped at the regional scale.

#### S1.2 Data

We used a combination of plot-level national forest inventory (NFI) data from the U.S. Forest Service Forest Inventory and Analysis (FIA) program and gridded datasets to train and run our model. We trained our biomass model at the FIA plot level, and then ran the model spatially using gridded datasets as inputs. When training the model at the plot level, we used FIA variables combined with PRISM meteorological data and mean NLCD canopy cover extracted in the 0.5-mile buffer around fuzzed FIA plot locations. We used the NLCD canopy cover rather than the FIA-recorded canopy cover because we use the gridded NLCD dataset to run our model spatially, and there are well-recorded discrepancies between plot-level canopy cover recorded by field crews and remote sensing based canopy cover products at the same location (70). When running the model spatially, we used corresponding gridded datasets for the predictor variables. Plot-level and gridded data sources for predictor variables are described in Table S1.

#### S1.2.1 National forest inventory data

We analyzed all 53,000 FIA plots in the Western United States that were more than 10% forest and were measured at least once since 2000. We use the FIA DataMart version 2.1.0, which uses new allometric models consistently across the U.S. (71). We aggregated tree-level biomass data and various condition-level variables to the plot level. We calculated plot-level biomass densities from tree-level biomass data, adjusting for the number of trees per acre that each sample tree represents, and the proportion of the plot that is in each condition. 23% of the forest plots consisted of more than one forested condition. We calculated plot-level estimates of continuous condition-level variables (canopy cover, stand age, slope, aspect, and elevation) as a weighted average based on the proportion of the plot that is in each condition. For categorical condition-level variables (ecosection, ecoprovince, ownership group type), we selected the category from whichever condition constituted the largest proportion of the plot. For each plot, we calculated the number of years since the fire occurred using FIA’s recorded disturbance codes. We considered a fire to have occurred if a fire disturbance code was recorded for any condition in that year. We defined fire as including all prescribed or natural crown and ground fires (disturbance codes 30, 31, and 32).

We included all 53,000 plots with forest cover greater than 10% when estimating initial biomass in approximately 2005 (Section 4.1.1). When estimating changes in biomass over time (Section 4.1.2) we further filtered the data to include only the 39,515 plots which were measured at least twice since 2000, and that remained greater than 10% forest across all measurements.

#### S1.2.2 Gridded data

We used a variety of gridded datasets to run our model (Table S1), and regridded all datasets to a consistent 1 km grid. We used monthly PRISM temperature, precipitation, and vapor pressure deficit data (26) to characterize the climate. We used annual tree canopy cover from NLCD (24). We used annual area burned from the MTBS-Interagency wildfire dataset (21) to quantify the number of years since fire occurred in each gridcell. We also used several static gridded variables: slope (72), aspect (73), elevation (74), ecosection and ecoprovince regions (75), and percent public ownership (76). After running our model spatially with gridded predictor datasets, we filtered the data to include only land classified as forest land use according to the Landscape Change Monitoring System (LCMS, 77), to be approximately consistent with the official USFS definition of forest land. We define forest land remaining forest land as areas categorized as forest land use for 20 or more consecutive years, following the U.S. Forest Service and Environmental Protection Agency approach (14).

#### S1.3 Model evaluation

We evaluated each random forest separately using the 20% of plots withheld from the training dataset for each ensemble member. Given our focus on biomass changes across the Western U.S. as a whole, our model is designed to capture large-scale spatial patterns rather than fine-scale spatial variability in biomass density.

We evaluate our model at the regional scale by coarsening our dataset to 0.25 degree resolution and averaging biomass estimates across the testing plots in those grid cells. At the regional scale, our model performs reasonably well. Our model can explain 94% of the variance in initial absolute biomass (Figure S11), and 54% of the variance in biomass changes between measurements (Figure S12). When considering each biomass change estimator separately, the fire (R²=0.56) and undisturbed (R²=0.53) model explain about the same fraction of variance at the grid cell level (Figure S12). We note that we include latitude and longitude as predictors for the initial biomass models, but not for the biomass change models, because the recursive nature of our model means that biases in initial biomass propagate to biomass change estimates.

Our model explains less of the variance in biomass when evaluated at the plot level, which is consistent with our expectations given the high level of variability at the plot level that our analysis is not attempting to capture. Furthermore, the fuzzed FIA plot locations prevent us from comparing our model estimates at the exact same location where plots were measured, and there is a high degree of small-scale spatial variability in biomass stock changes. Our model can explain 68% of the variance in initial absolute biomass at the plot level, and 55% of the variance in biomass changes at the plot level (Figure S11). Our estimators for carbon changes following fire have greater predictive power than our estimators for carbon changes in unburned plots. The fire carbon change model explains 55% of plot-level variance, respectively, compared to only 24% for the undisturbed estimator. The poorer performance at the plot level is expected because there are high levels of variability at smaller spatial scales that we are not attempting to capture in our approach.

#### S1.4 Open source tools

All analyses were performed using Python open source packages (Cartopy (78), Geopandas (79), Joblib (80), Jupyter (81), Matplotlib (82), NumPy (83), Pandas (84, 85), Rasterio (86), Rioxarray (87), Shap (68), Shapely (88), Sklearn (89), Pint (90), Pyproj (91), and Xarray (92)).

### S2 Supplementary figures

**Figure S1:**
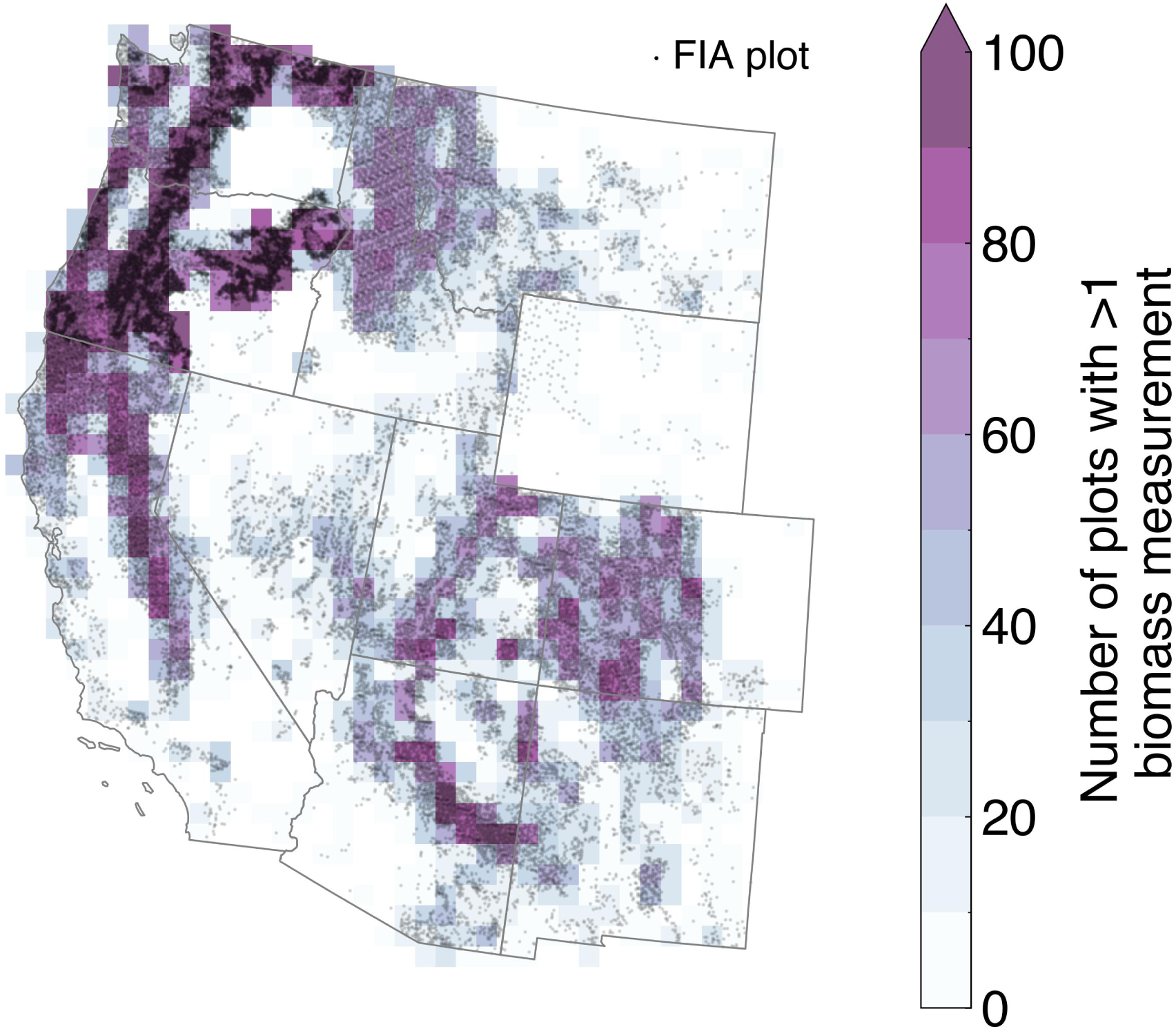
Forest inventory plots analyzed in this study. Gridded colors indicate the number of forest inventory plots with at least two recorded biomass measurements in each 0.25 degree grid cell. Gray points indicate the fuzzed location of the 39,515 forest inventory plots in the Western United States with a forest condition and at least two recorded biomass measurements.

**Figure S2:**
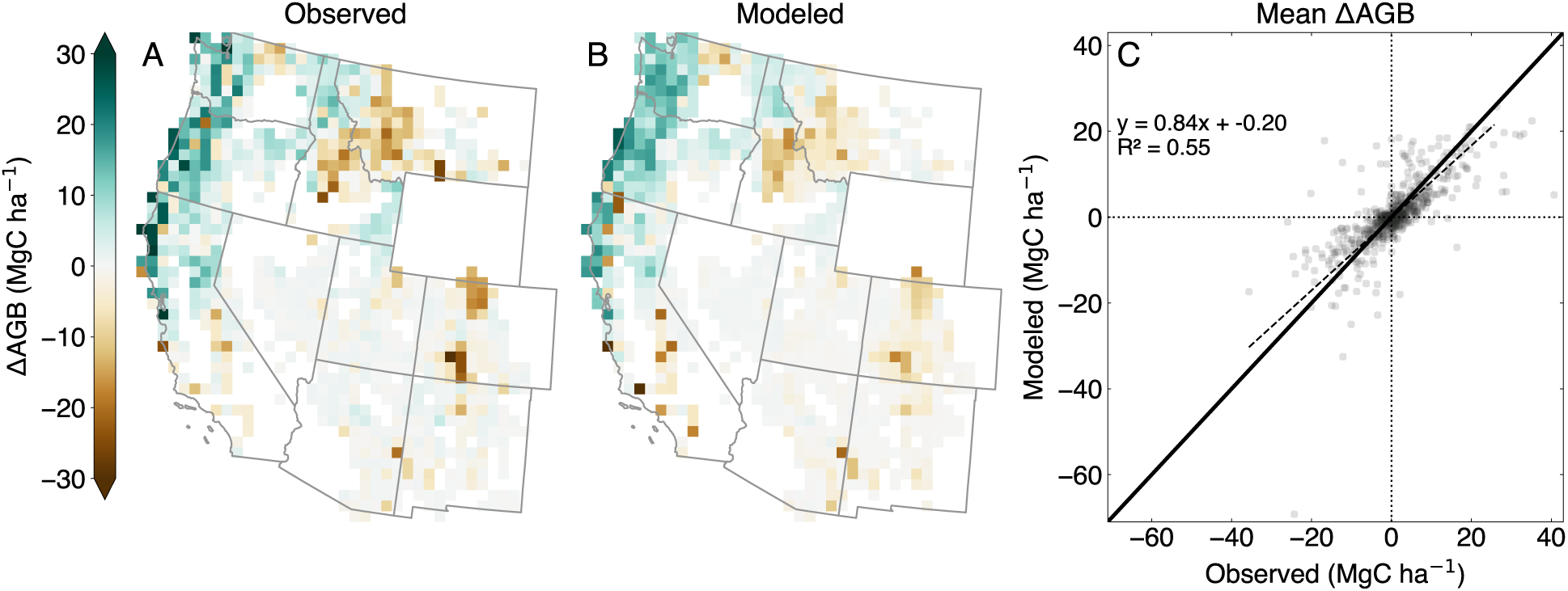
Comparison to forest inventory measurements. Spatial distribution of plot-level changes in live aboveground biomass (AGB), averaged across 0.25 degree grid cells, for FIA observations (A) and our model estimates (B). In (B), modeled biomass estimates are extracted in the 0.5 mile (0.8 km) buffer surrounding the fuzzed FIA plot locations in the same years that the plots were measured in (A). Only grid cells containing more than 10 remeasured FIA plots are shown. No grid cells in Wyoming meet that criteria due to Wyoming’s late entry into the current FIA inventory protocol (14). (C) Comparison of grid cell averages in (A) vs. grid cell averages in (B). The 1:1 line is shown in solid black, and the ordinary least squares regression fit is shown in dashed black.

**Figure S3:**
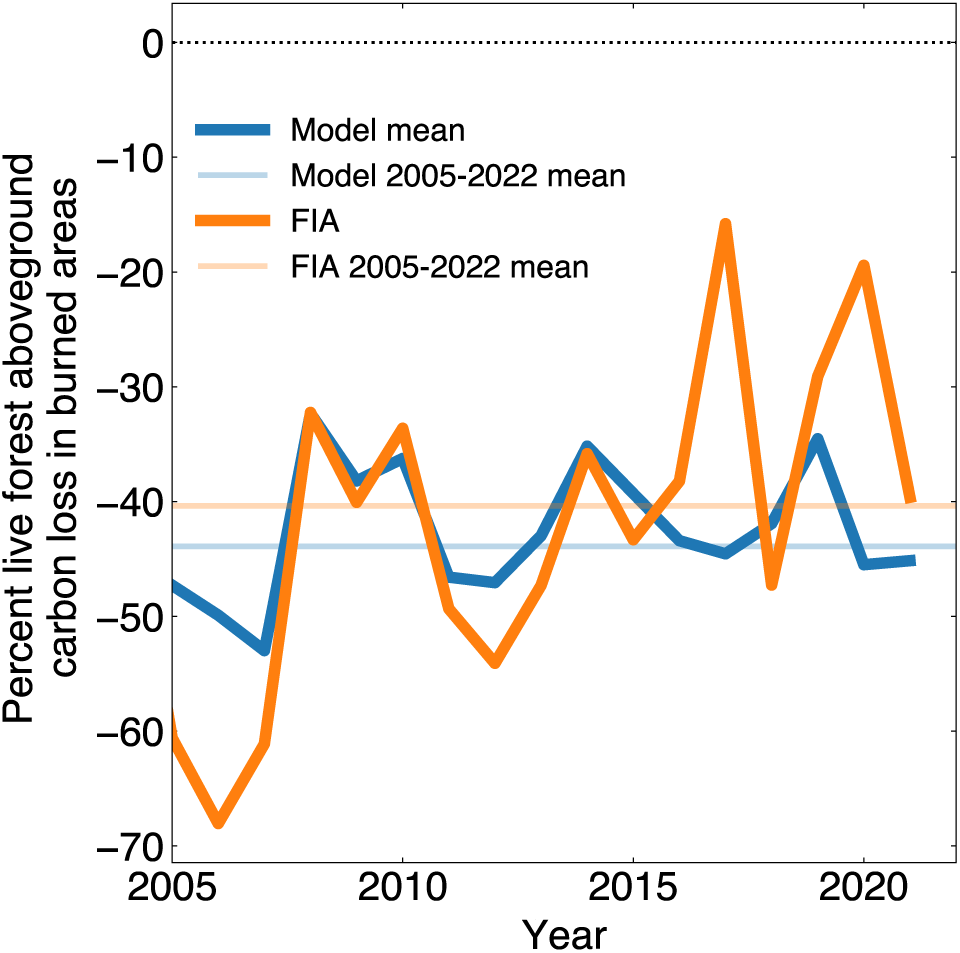
Mean percent change in live aboveground carbon in burned plots from 2005 to 2022, in our model (blue) and in FIA data (orange). Modeled live biomass carbon losses from tree mortality is 43% on average, and approximately constant over time and consistent with the mean 40% carbon losses from tree mortality rates observed in FIA plots.

**Figure S4:**
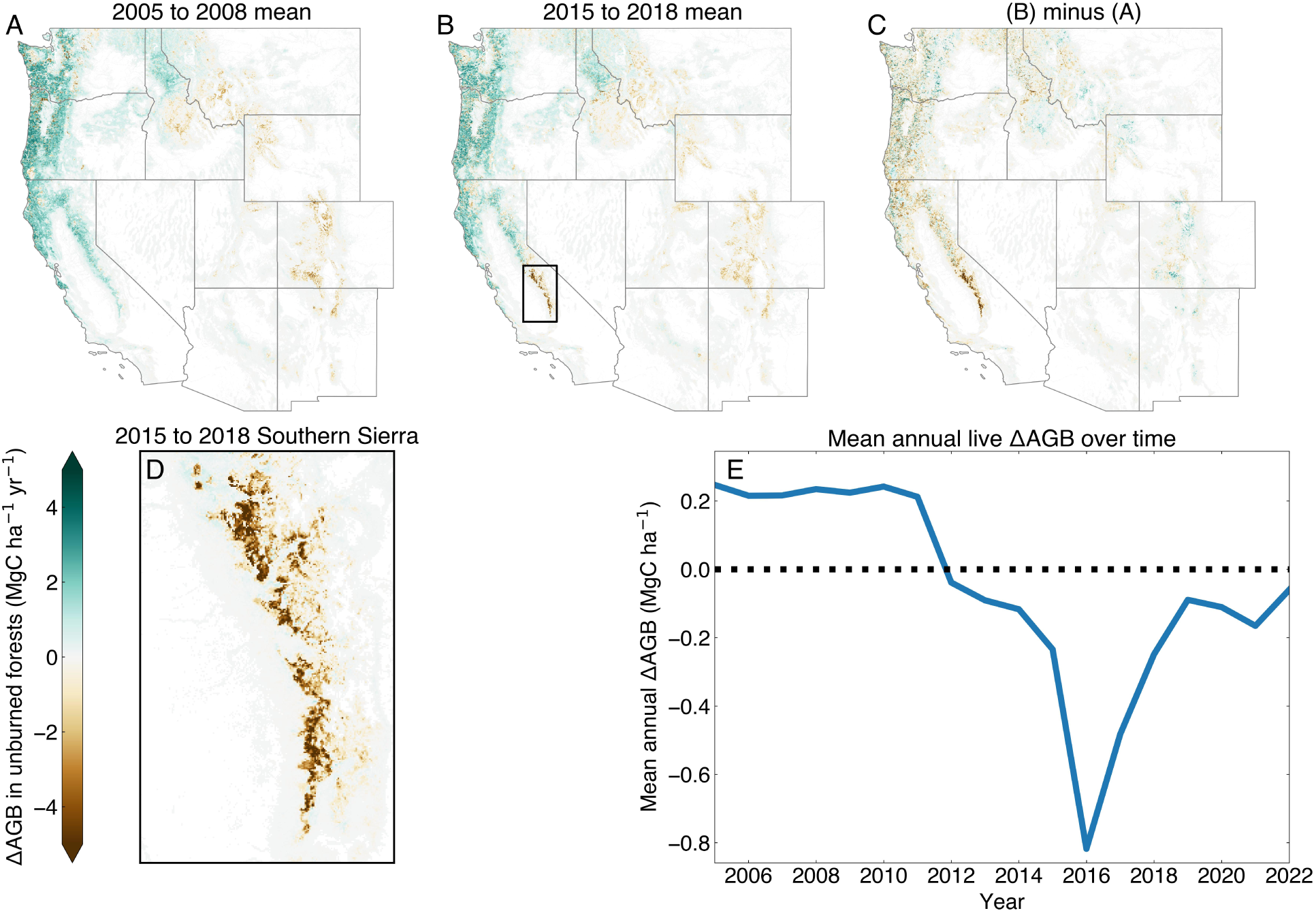
Comparison of changes in annual mean change in live aboveground biomass (AGB) in unburned forests during (A) 2005 to 2008 and (B) 2015 to 2018; and (C) the difference in annual mean change in live AGB in unburned forests between these two time periods (B) - (A). The mean live AGB change in unburned forests from (B) is shown in the Southern Sierra in (D). (E) shows the time series of change in live AGB in unburned forests across the Southern Sierra, averaged across the spatial domain shown in (D).

**Figure S5:**
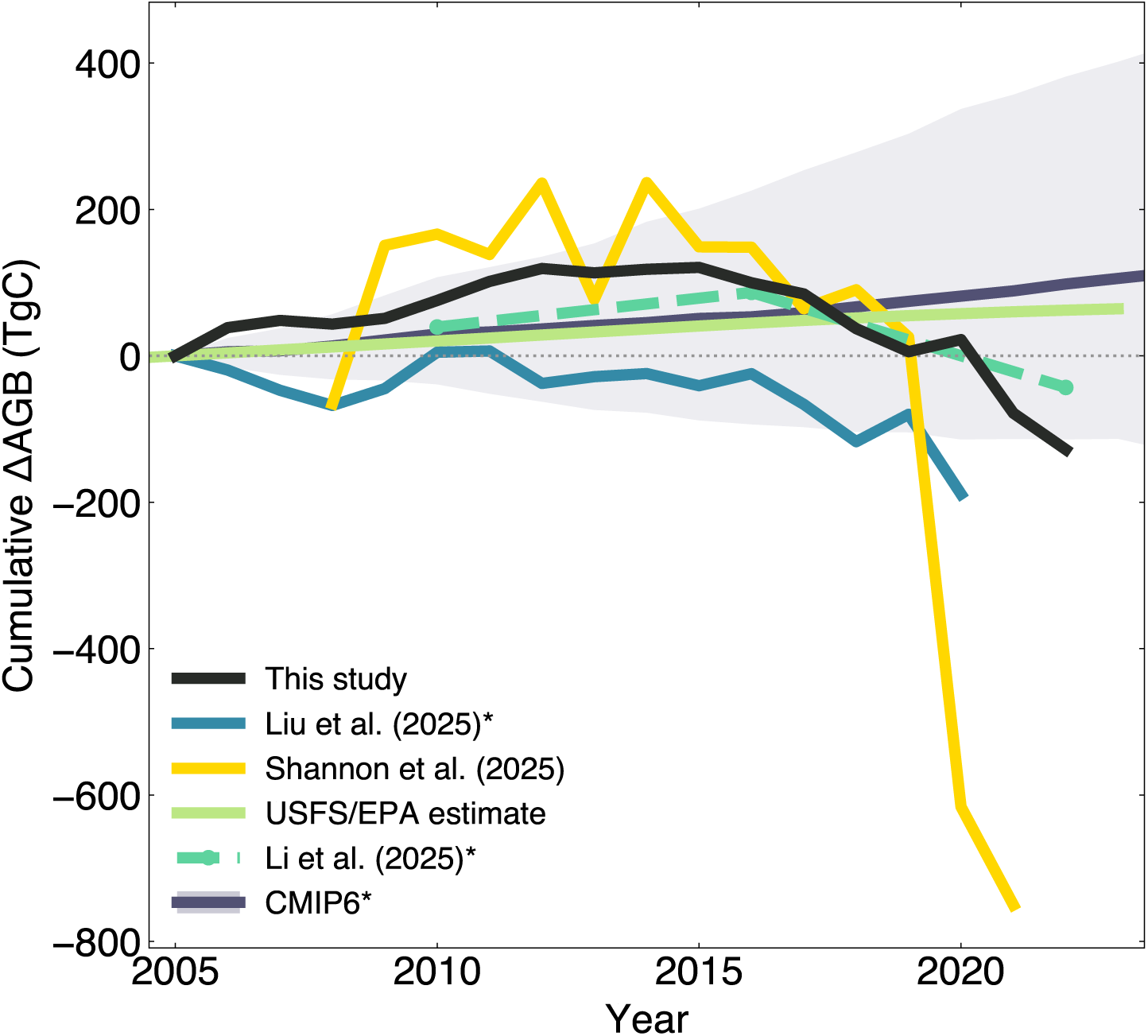
Comparison to trends in other biomass products (as in Figure 4, but where starred datasets (Li et al. 2025, Liu et al. 2025, CMIP6) are normalized by the percent of forest remaining forest.

**Figure S6:**
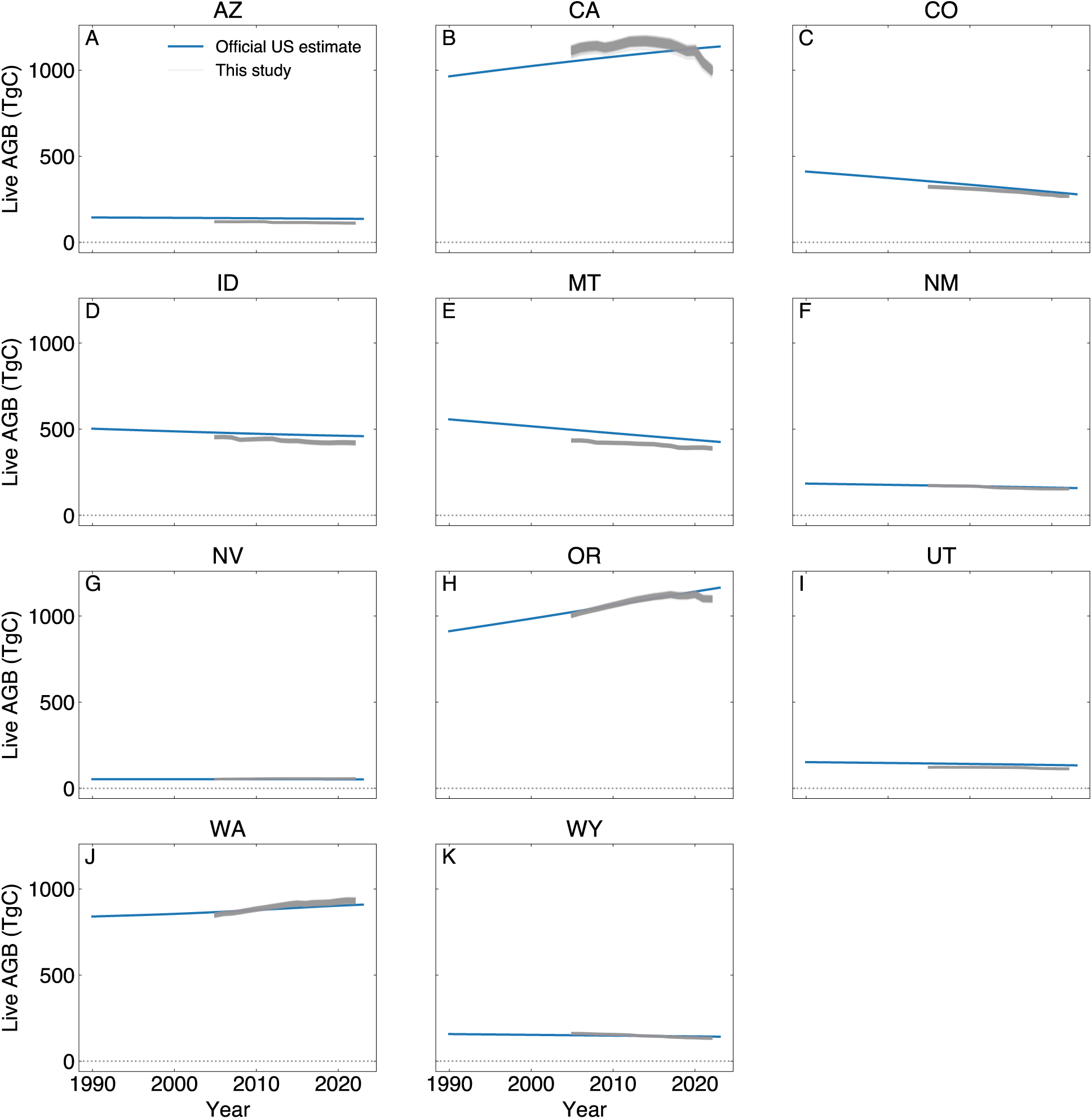
Forest live aboveground biomass over time (as in Figure 2A) for individual Western states. Each gray line is a different ensemble member. All states share the same y-axis scale.

**Figure S7:**
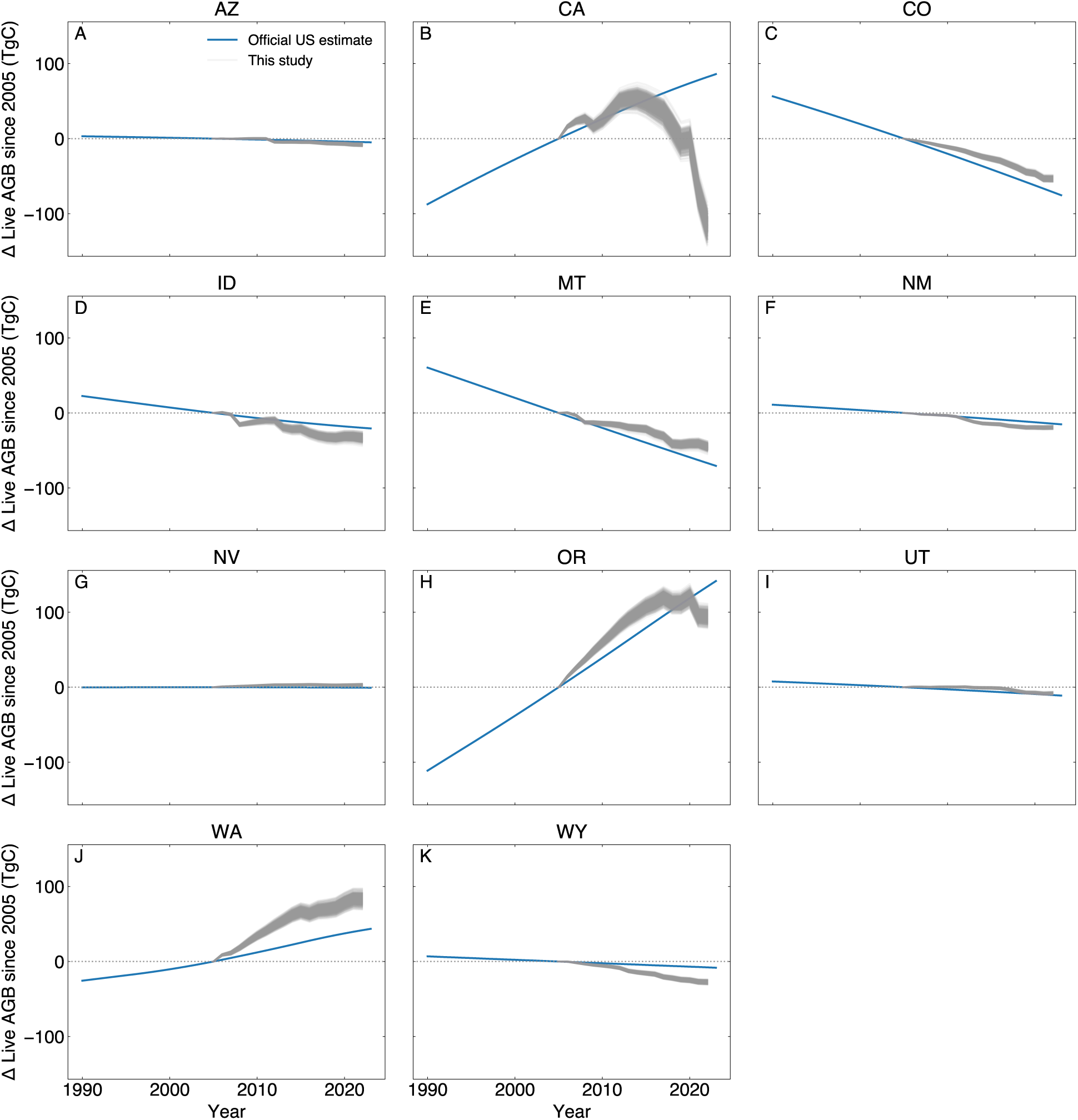
Change in forest live aboveground biomass over time for individual Western states. Each gray line is a different ensemble member. All states share the same y-axis scale.

**Figure S8:**
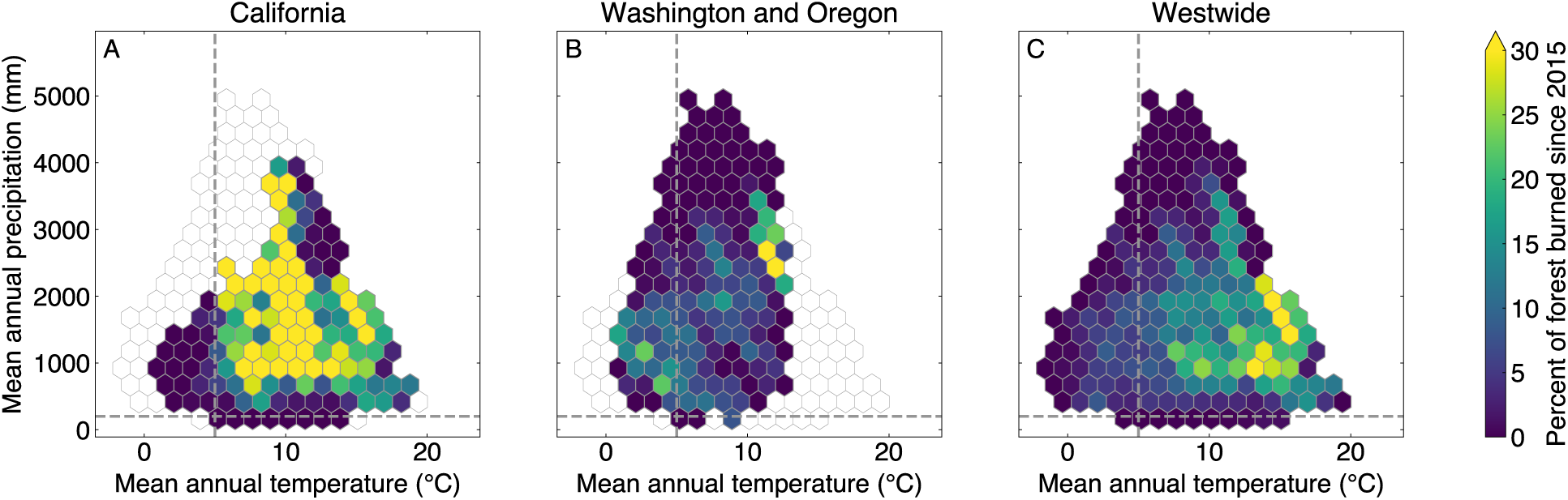
Percent of forest area burned from 2015 to 2022, plotted across climate space for (A) California, (B) Washington and Oregon, and (C) the Western United States. Hexagonal bins are plotted when the bin contains at least 25 1,000 meter grid cells that contain more than 50% forest. The horizontal and vertical dashed lines represent an approximate climate threshold for defining “arid forests” (precipitation <150 mm/year, mean annual temperature >5°C). 23% of arid forests burned in California from 2015 to 2022, compared to 7% in Washington and Oregon.

**Figure S9:**
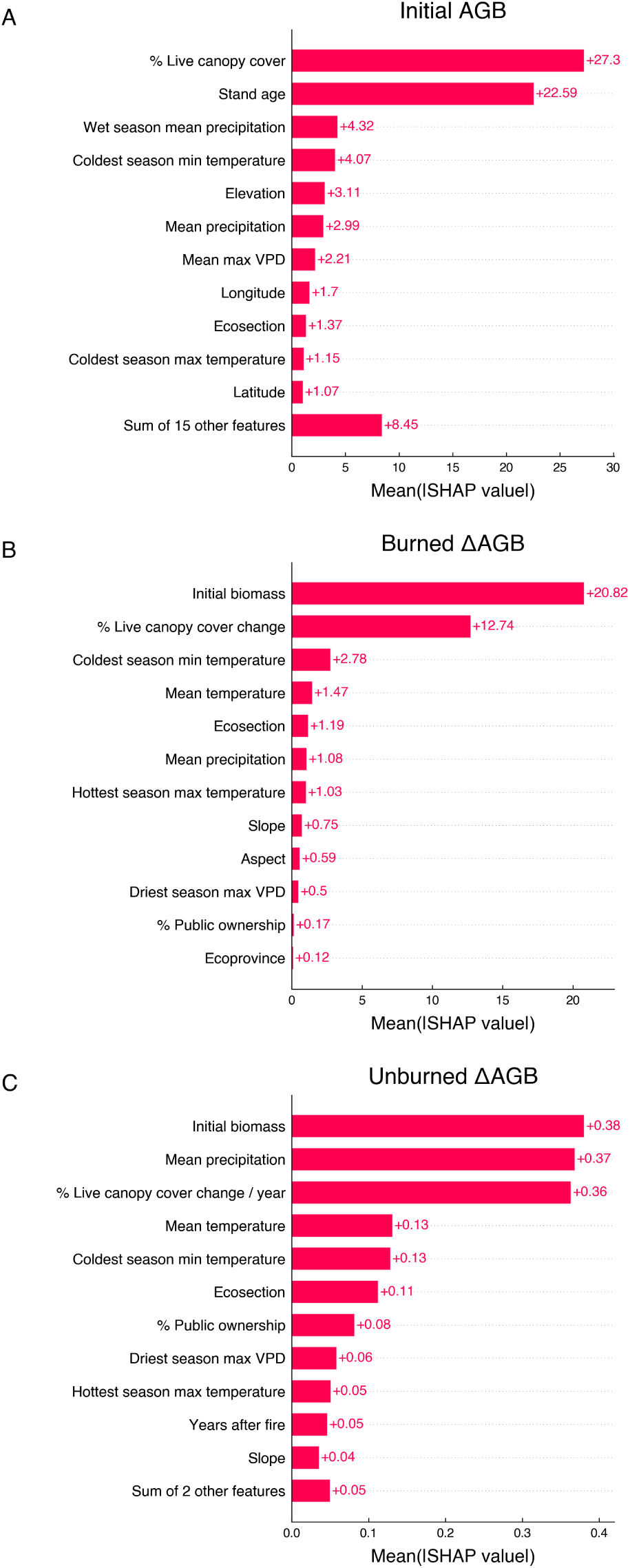
Feature importance of random forest models constructed for (A) initial live aboveground biomass (AGB), (B) live AGB change in burned forests, and (C) live AGB change in unburned forests.

**Figure S10:**
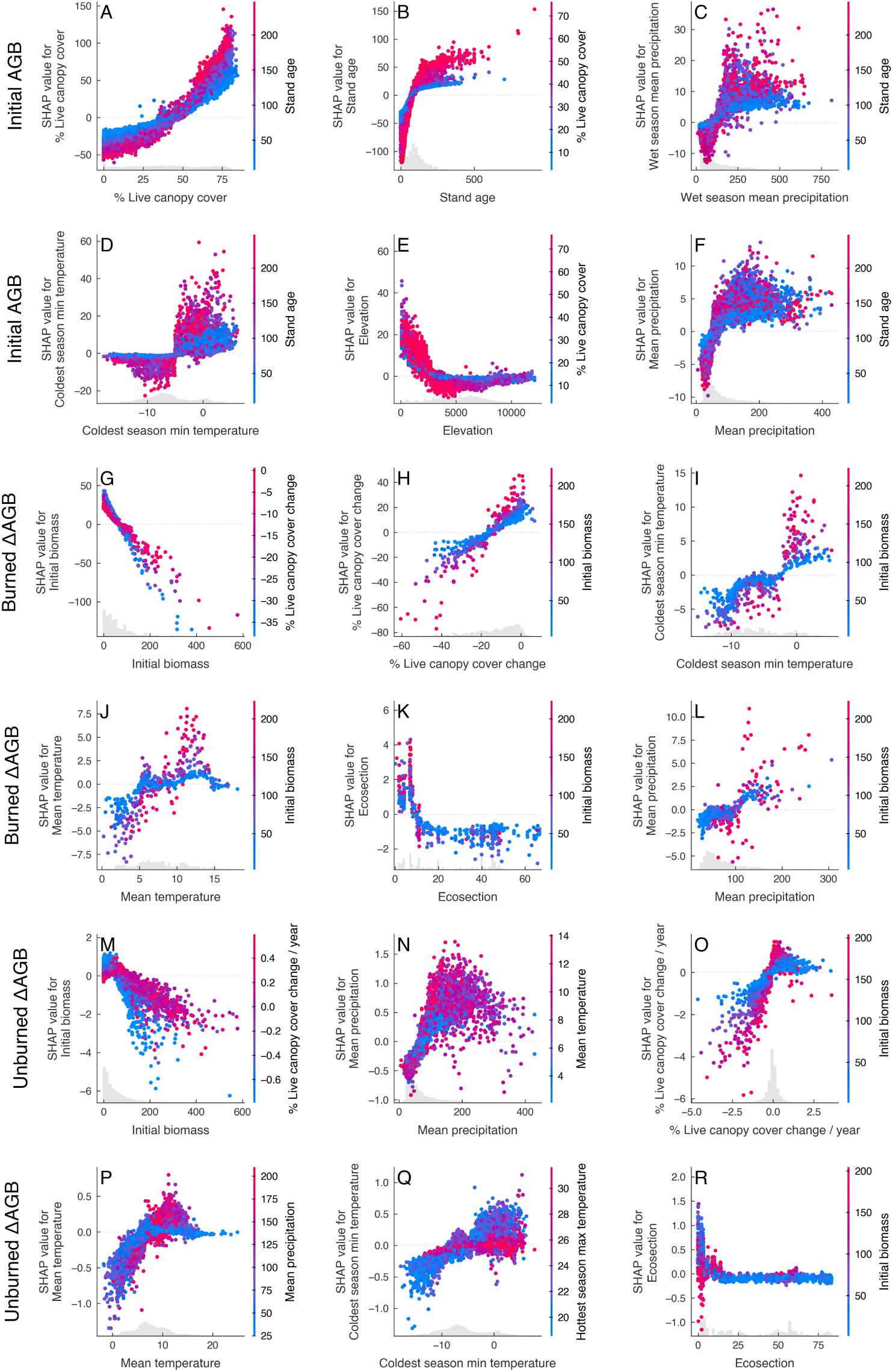
Partial dependency plots for the six most important features (see Figure S9) for the initial live aboveground biomass (AGB) model (A-F), the model live AGB change in burned forests (G-L), and the model of live AGB change in unburned forests (M-R). Each point represents one plot, the x-axis indicates the feature value, and the y-axis indicates that feature’s SHapley Additive exPlanation (SHAP) value. SHAP values above zero mean that the feature increased that plot’s prediction relative to a baseline, and SHAP values below zero mean that the feature decreased that plot’s prediction. Color indicates the feature with the strongest interaction with the x-axis feature.

**Figure S11:**
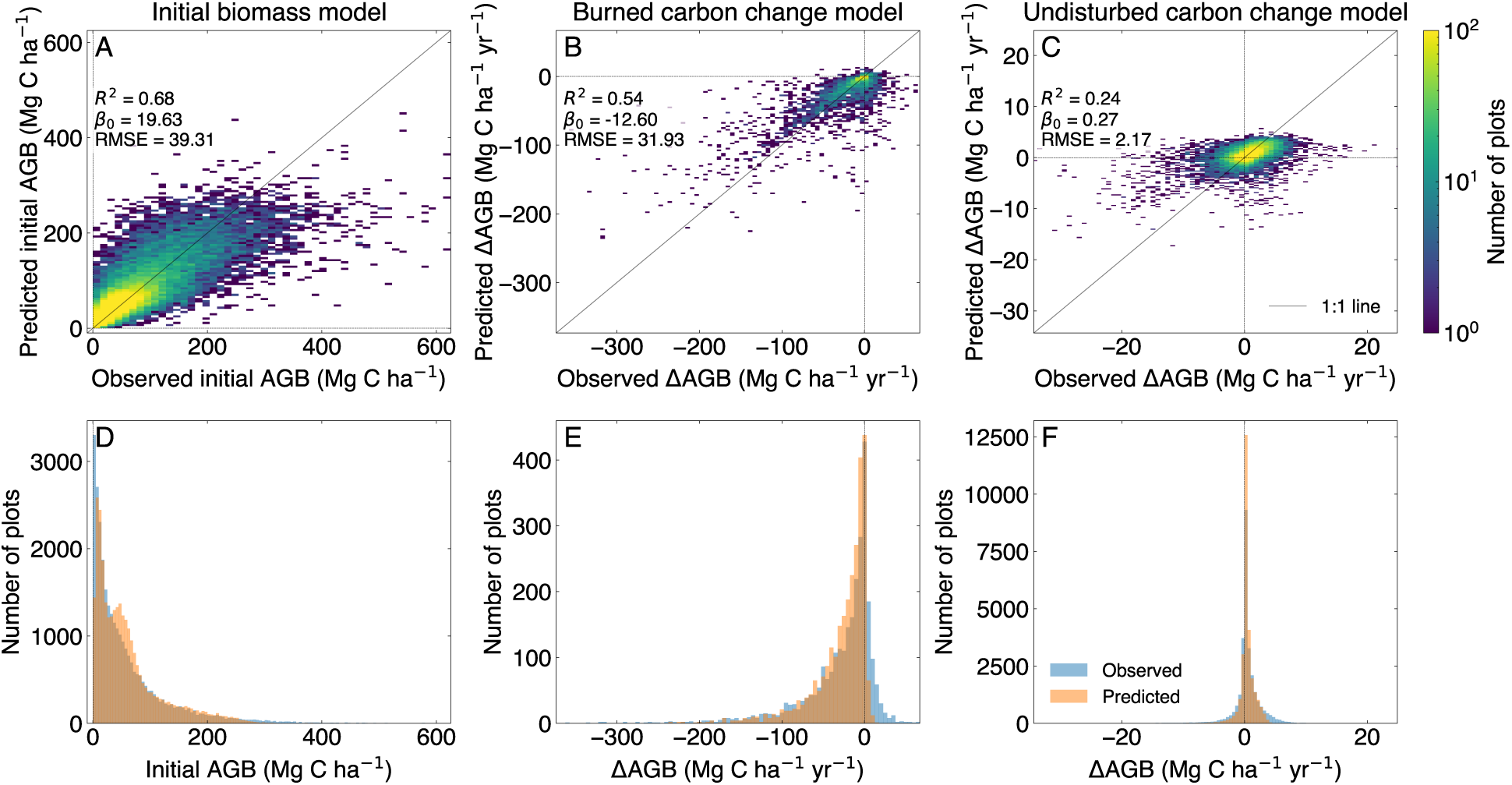
Plot-level evaluation of live aboveground biomass (AGB) model. Observed vs. predicted initial live AGB (A), change in AGB in burned forests (B), and change in AGB in unburned forests (C). Color indicates the density of plots in each 2D histogram grid cell, on a log scale. Histograms of the distributions of observed and predicted initial live AGB (D), change in AGB in burned forests (E), and change in AGB in unburned forests (F). All panels show predictions for test plots withheld from model training. For plots withheld by more than one ensemble member, the mean predicted value across ensemble members is shown.

**Figure S12:**
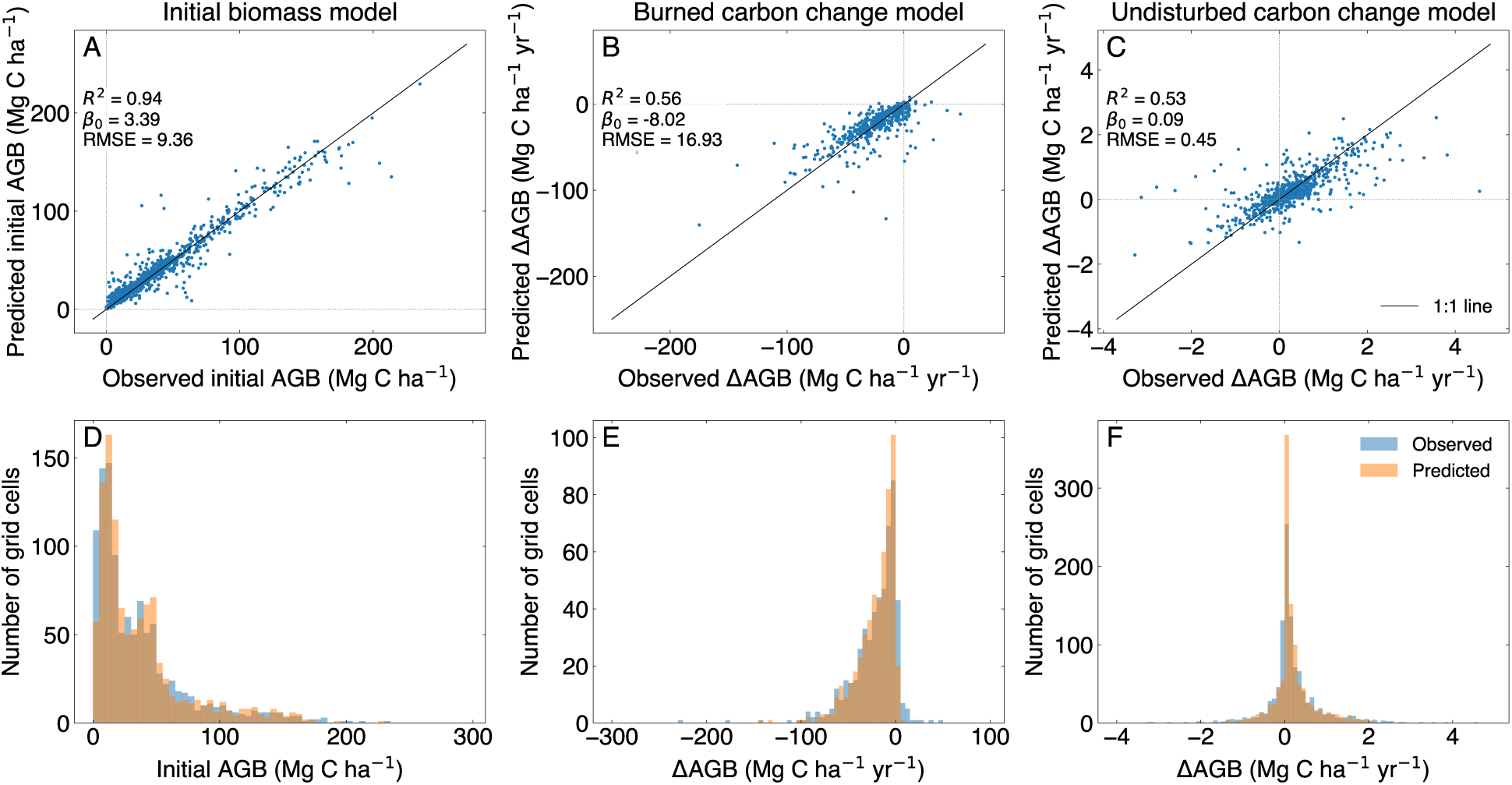
Evaluation of live aboveground biomass model, where plot-level data is averaged into 0.25 by 0.25 degree grid cells, using the grid shown in Figure S2. Observed vs. predicted gridcell-mean initial live AGB (A), change in gridcell-mean AGB in burned forests (B), and change in gridcell-mean AGB in unburned forests (C). Histograms of the distributions of observed and predicted initial live AGB (D), change in AGB in burned forests (E), and change in AGB in unburned forests (F). All panels show grid cell average of predictions for test plots withheld from model training.

### S3 Supplementary tables

**Table S1:**
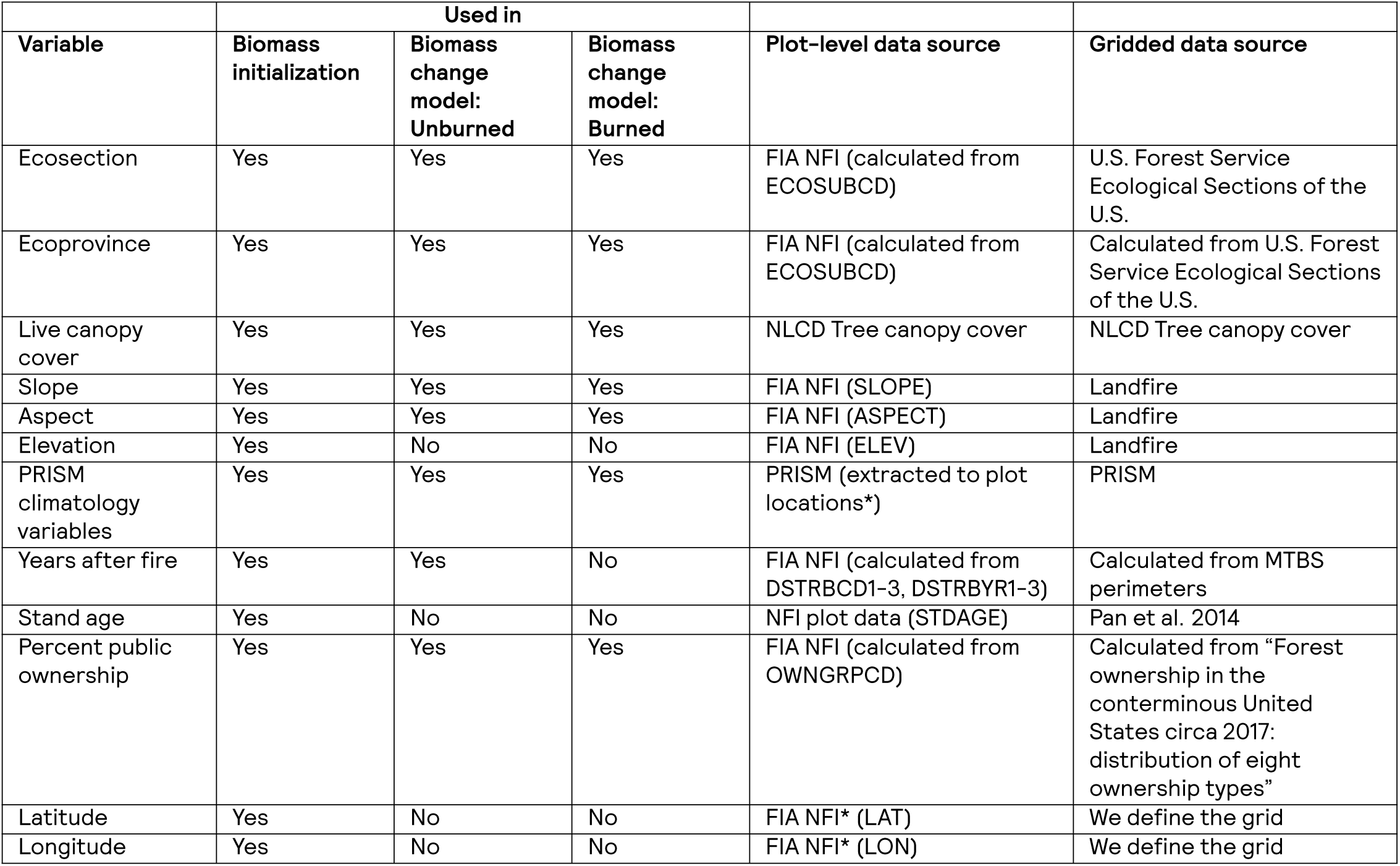
Variables used in model, and sources for plot-level and gridded data. FIA NFI refers to plot-level data from the Forest Inventory and Analysis Program National Forest Inventory (23). * Note that the coordinates of FIA plots are fuzzed and swapped. We used them as if they were exact.

**Table S2:** Fossil fuel CO₂ emissions from the U.S. Environmental Protection Agency state-level greenhouse gas inventory (93).

| State | CO<sub>2</sub> fossil emissions (TgC) |
| --- | --- |
| Arizona | 191.8 |
| California | 709.7 |
| Colorado | 190.2 |
| Idaho | 40.0 |
| Montana | 65.0 |
| New Mexico | 104.2 |
| Nevada | 84.0 |
| Oregon | 82.9 |
| Utah | 130.4 |
| Washington | 150.8 |
| Wyoming | 128.2 |
| **Total** | **1877.2** |

**Table S3:** Percent of forest area experiencing net gain vs. net loss of live biomass.

|  | Percent of forest area |  |  |
| --- | --- | --- | --- |
|  | Net gain (> +1 MgC ha<sup>-1</sup>) | Net loss (< -1 MgC ha<sup>-1</sup>) | Minimal change (-1 to +1 MgC ha<sup>-1</sup>) |
| 2005 to 2015 | 31% | 15% | 54% |
| 2015 to 2022 | 19% | 18% | 63% |
| 2005 to 2022 | 32% | 22% | 46% |

**Table S4:** Rates of live biomass change in burned vs. unburned locations, for different time periods.

|  | Mean annual Western U.S. live biomass carbon flux (TgC yr<sup>-1</sup>) |  |  |
| --- | --- | --- | --- |
|  | Fire losses | Unburned change | Net change |
| 2005 to 2015 | -16.7 | 28.0 | 11.3 |
| 2015 to 2022 | -45.8 | 19.2 | -26.6 |
| Change between time periods | -29.1 (76.8%) | -8.8 (23.2%) | -37.9 |

**Table S5:** Number of ensemble members with cVeg data available for both the historical and SSP245 experiments, per CMIP6 model.

| Model | Number of ensemble members |
| --- | --- |
| ACCESS-ESM1-5 | 40 |
| CanESM5 | 50 |
| CanESM5-1 | 20 |
| CanESM5-CanOE | 3 |
| MIROC-ES2L | 28 |
| MIROC-ES2H | 3 |
| IPSL-CM6A-LR | 11 |
| CNRM-ESM2-1 | 9 |
| CMCC-CM2-SR5 | 1 |
| CMCC-ESM2 | 1 |
| BCC-CSM2-MR | 1 |
| KIOST-ESM | 1 |
| TaiESM1 | 1 |
| **Total** | **169** |

## Notes

https://doi.org/10.5281/zenodo.19698817

